# Misexpression of the Homologues of Bryophyte Gametophyte-to-Sporophyte Control Genes in Arabidopsis Results in Germline Reprogramming and Phenotypes that Mirror Apomictic Development

**DOI:** 10.1101/2022.09.15.508103

**Authors:** William Bezodis, Helen Prescott, Hugh Dickinson

## Abstract

Evidence from the model bryophytes *Physcomitrium* and *Marchantia* suggests that a BELL-KNOX genetic module acts as a master regulator controlling sporophyte identity. Investigating any conservation of this system in flowering plants has proved challenging, but studies of the *Arabidopsis eostre* mutant and naturally apomictic angiosperms point to ectopic activation of KNOX and BELL transcription factors mediating the switch from sexual to apomictic development. We show here that in *Arabidopsis*, ectopic expression, under a germline-specific promoter, of KNOX and BELL genes not normally expressed in the gametophytes both disrupts germ cell specification and causes defects in cell identity throughout gametophyte development – some mirroring events seen in naturally apomictic plants. A better understanding of this TALE-HD genetic module in flowering plants may thus help to unravel the molecular control of higher plant life cycles, while providing a route to engineering synthetic apomixis in crops. This study also highlights the utility of applying data from bryophytes, where the ontogeny transitions are spatio-temporally distinct, to apomixis research in angiosperms.

## Background

Alternation of generations is fundamental to the land plant life cycle and involves switching between multicellular haploid gametophyte and diploid sporophyte ontogenies. In flowering plants such as *Arabidopsis*, the gametophyte has been reduced from the dominant photosynthetic stage to 3-7 cells nutritionally dependent on the sporophyte (Friedman, 2013). The molecular control of alternation has been studied in the moss *Physcomitrium patens* (summarised by Horst and Reski, 2016) where TALE-HD transcription factors of the KNOX and BELL classes are thought to regulate the switch from gametophyte to sporophyte by promoting the sporophyte ontogeny, being repressed in the gametophyte by PRC2-mediated H3K27me3 histone modification (Bowman *et al*., 2016). Horst *et al*. (2016) identified *PpBELL1* as a master sporophyte regulator, showing that misexpression in the gametophyte could induce a switch to sporophyte identity with largely functional ectopic sporophytes produced. Further, Sakakibara *et al*. (2013) showed that deletion of both *Physcomitrium KNOX2* genes prevented sporophyte development, such that *KNOX2* knockout results in plants constitutively adopting the gametophyte ontogeny, the opposite of *PpBELL1* overexpression. BELL and KNOX transcription factors function by heterodimerisation, therefore pointing to a dual role of a KNOX+BELL module. However, questions remain about whether (i) this module functions in the gamete in addition to the sporophyte and (ii) the extent to which this function is conserved, as it may differ between *Physcomitrium* and *Marchantia* (Horst *et al*., 2016; Hisanaga *et al*., 2021). It has also been proposed that BELL/KNOX control gamete identity, instead of or in addition to sporophyte identity (Ortiz-Ramirez *et al*., 2017; Dierschke *et al*., 2021; Hisanaga *et al*., 2021), but it is unclear how these putative roles interact.

This genetic control in bryophytes may have evolved, along with the appearance of the sporophyte, from an ancestral algal system controlling the diploid zygote (Bowman *et al*., 2016), such as is present in extant *Chlamydomonas reinhardtii*. The plus and minus gametes of *Chlamydomonas* express a BELL and KNOX respectively, and ectopic expression is sufficient to activate zygote development in vegetative cells (Lee *et al*., 2008). It has been suggested that this genetic system may be conserved across land plants (Horst and Reski, 2016; Bowman *et al.*, 2016) including in all crop species, but convincing data are lacking.

In angiosperms, any role for BELL-KNOX as a regulator of alternation of generations is obscured by the increased number and functional diversity of these genes, which play varied roles in sporophyte development (Hay and Tsiantis, 2010). Nonetheless, there is evidence from *Arabidopsis* mutants that representatives of both the BELL (*BLH1*) and KNOX2 (*KNAT*) gene families may carry out such a conserved alternation function. For example, *BLH1* is not expressed in WT gametophytes but is expressed ectopically in the *eostre* mutant (Pagnussat *et al*., 2007), causing developmental defects including aborted embryo sacs, abnormal nuclear migration and fate switch from synergid to egg cell, indicative of both sporophyte and gamete control functions. Furthermore, *BLH1* interacts with *KNAT3* (Hackbusch *et al*., 2005) and crossing *blh1 knat3* knockout lines into *eostre* mutants resulted in a partial suppression of the phenotype (Pagnussat *et al*., 2007). However, as *KNAT3* is not reported to be expressed in the gametophyte, the mechanism underlying this effect remains unclear. Pagnussat *et al*., (2007) also showed that the *OVATE FAMILY PROTEIN5* (*OFP5*) knockout phenocopies the *eostre* mutant. Several *OFP* proteins in this clade have been shown to interact with *BLH1* and *KNATs* and regulate their cellular localisation (Hackbusch *et al*., 2005; Wang *et al*., 2016).

Interestingly, naturally apomictic mechanisms in some angiosperms appear to involve *BLH1* misexpression in the female gametophyte, reminiscent of the *eostre* mutant (Pagnussat *et al*., 2007; Okada *et al*., 2013; Galla *et al*., 2019). In *Hieracium praealtum*, the *BLH1* orthologue is expressed in apomictic aposporous initial cells and early embryosac cells (Okada *et al*., 2013) accompanied by phenotypic similarities between apomictic *Hieracium* embryo sacs and the *eostre* mutant. Likewise, the facultative apomict *Hypericum perforatum* also expresses the *BLH1* orthologue in apomictic, but not sexual gametophytes (Galla *et al*., 2019). This may reflect similar roles for BLH1 in inducing the *eostre* phenotype, regulating apomictic development, and by analogy with events in bryophytes, alternation of generation. This research illustrates the utility of applying principles derived from bryophyte developmental genetics to angiosperm research. As the sporophyte and gametophyte are more distinct in bryophytes, it is easier to untangle reproductive transitions in these models which can then be translated to angiosperm models like *Arabidopsis* and, in future, potentially into crop species.

Previous exploration of the roles of TALE-HD genes in flowering plant alternation using knockout lines have been confounded by the impact of these mutations on vegetative development, making accurate determination of any specific effect on reproductive cell linages difficult (Furumizu *et al*., 2016). Here, we report inducible ectopic expression, under a germline promoter, of *BLH1*, and 4 *KNOX2* genes not normally expressed in male and female gametophytes. Not only do these lines recapitulate the *eostre* mutant phenotype, but germ cell specification is also reprogrammed, with defects to cell identities occurring throughout gametophyte development. Importantly, as with the *eostre* mutant, some of the phenotypes observed mirror events seen in naturally apomictic plants.

## Results

### promSPOROCYTELESS Drives TALE-HD Expression in the Male and Female Germline

To confine ectopic expression of BELL and KNOX2 genes to male and female reproductive cell linages, the *SPOROCYTELESS* (SPL) promoter was employed in Golden Gate constructs to drive the LhGR element of a two-component inducible expression system (Craft *et al*., 2005). Treatment of plants with dexamethasone (Dex) thus activated pOp6 promotors driving expression of the TALE HD genes just prior to the establishment of gametophytic fate, in the tapetum, and throughout male and female gametophyte development (Fig1, in Mendes *et al*, 2020). These constructs were used to make plants containing all four *Arabidopsis* Class 2 KNOX genes not expressed in gametophytes (*KNAT3,4,5*, and *7*, henceforth referred to as ‘Quad*KNAT’* lines) by crossing two pairs of ‘TwinK*NAT*’ lines expressing *KNAT3&4*, and *KNAT5&7*. A further construct that expressed *BLH1* was produced and combined with *KNATs* through crossing. The constructs were transformed into both Col-0 *Arabidopsis* and a verified homozygous *ofp1345* knockout line (kindly provided by Zhang *et al*., 2020). The latter is the complete *OFP5-*clade, containing OFPs known to interact with *BLH1* and KNOX2 (Hackbusch *et al*., 2005). Plants expressing these constructs were validated for both transgene expression level and localisation (Figure 1).

**Figure 1.**
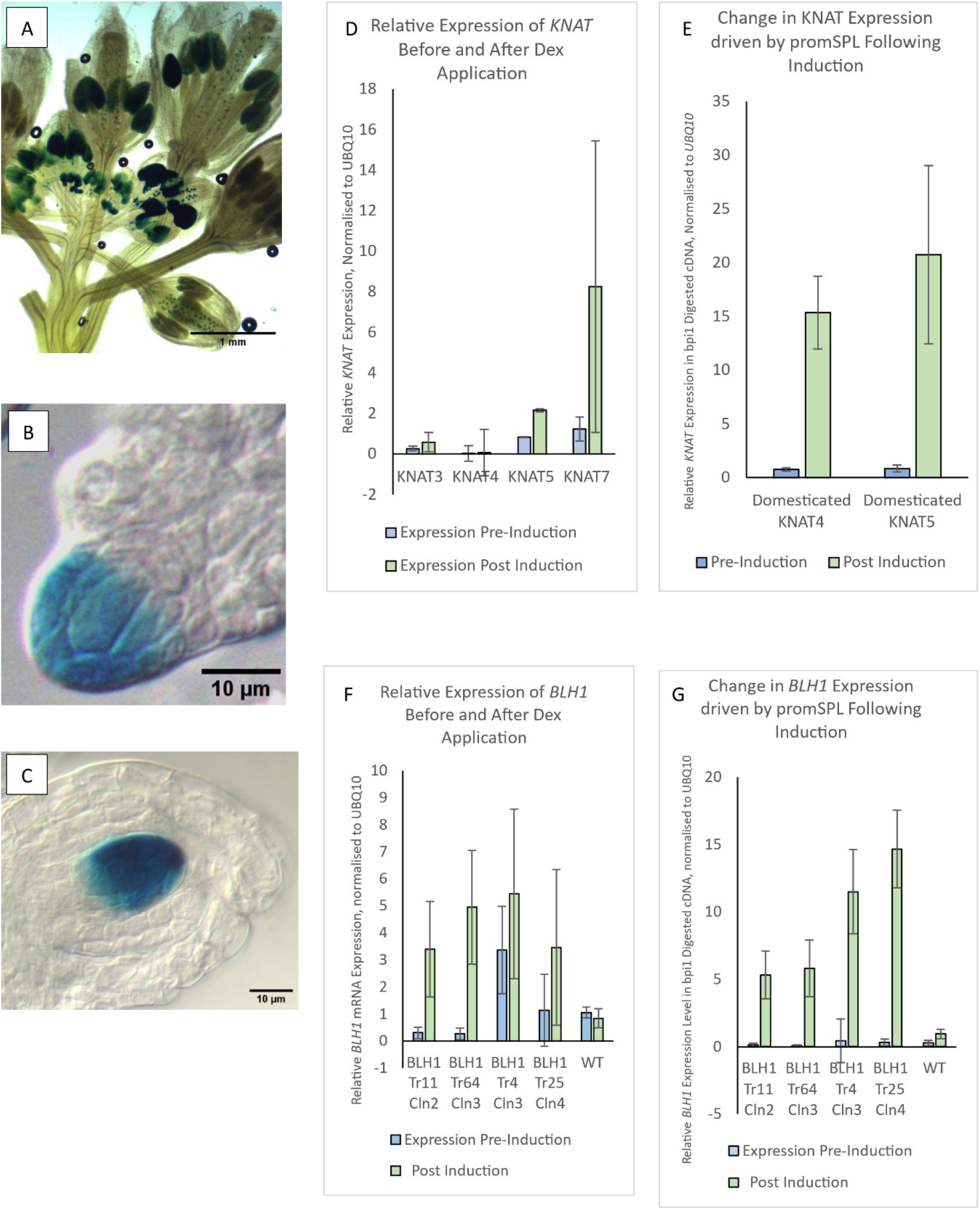
Dexamethasone induced *promSPOROCYTELESS* driven transgene expression pattern, and level of gene expression measured by RT-qPCR. **(A)** - **(C)** expression domain of a *promSPOROCYTELESS:GUS* reporter construct. **(A)** expression in anthers at different stages of development, **(B)** pre-meiotic female expression, and **(C)** post-meiotic female expression. **(D)** - **(G)** RT-qPCR data for Class 2 KNOX and *BLH1* from dexamethasone-induced inflorescences. **(D)** and **(E)** class 2 KNOX, **(F)** and **(G)** independent transformants of BLH1. Trn=transformant, cln=clone **(D)** and **(F)** show total cDNA, **(E)** and **(G)** show cDNA digested with bpi1, and using primers adjacent to an excised bpi1 restriction site to amplify only the ‘domesticated’ form of the gene. i.e. that has been synthesised without the bpi1 site and introduced as a transgene.

### Misexpression of TALE-HD Transcription Factors Causes the Specification of Multiple Functioning Germlines

A common defect in *Arabidopsis* early female development is the specification of multiple Megaspore Mother Cells (MMCs). This phenotype is present at a low level in WT but occurs frequently in epigenetic mutants (Rodriguez-leal *et al*., 2015; Mendes *et al*., 2020), and mutants with cell-cell communication defects (Su *et al*., 2020; Cai *et al*., 2022). This phenotype occurred extremely frequently in several of our experimental lines (fig2), particularly in Quad*KNAT* and *KNAT5*,7, often with 4-5 MMC-like cells within a single ovule in up to 100% of ovules in some gynoecia. These MMC-like cells acquire MMC identity, as confirmed by the MMC-specific KNUCKLES-YFP reporter (Payne *et al*., 2004; Tucker *et al*., 2012) (fig2). The effect of *promSPL:BLH1* expression on pre-meiotic MMC specification is relatively minor, but post-meiosis, Functional Megaspore Cell (FMC) development was frequently disrupted, with excess callose deposited. In both *promSPL:BLH1* and *SPL:KNAT* ovules with multi-MMCs, often more than one MMC appeared meiotically functional, resulting in dual/multiple gametophytes that arrest by the FG2 stage. Again, callose staining confirmed the presence of multi-meiotic nascent gametophytes (Zhao *et al*., 2014; Cai *et al*., 2022) with excessive callose deposition in an atypical pattern, often present in multiple cells within the nucellus. This phenotype resembles certain types of apomictic development (Tucker *et. al*, 2001; Galla *et. al*, 2011) and has previously been used as an indicator of a reported multiple fully functional MMC phenotype (Cai *et al*., 2022).

**Figure 2.**
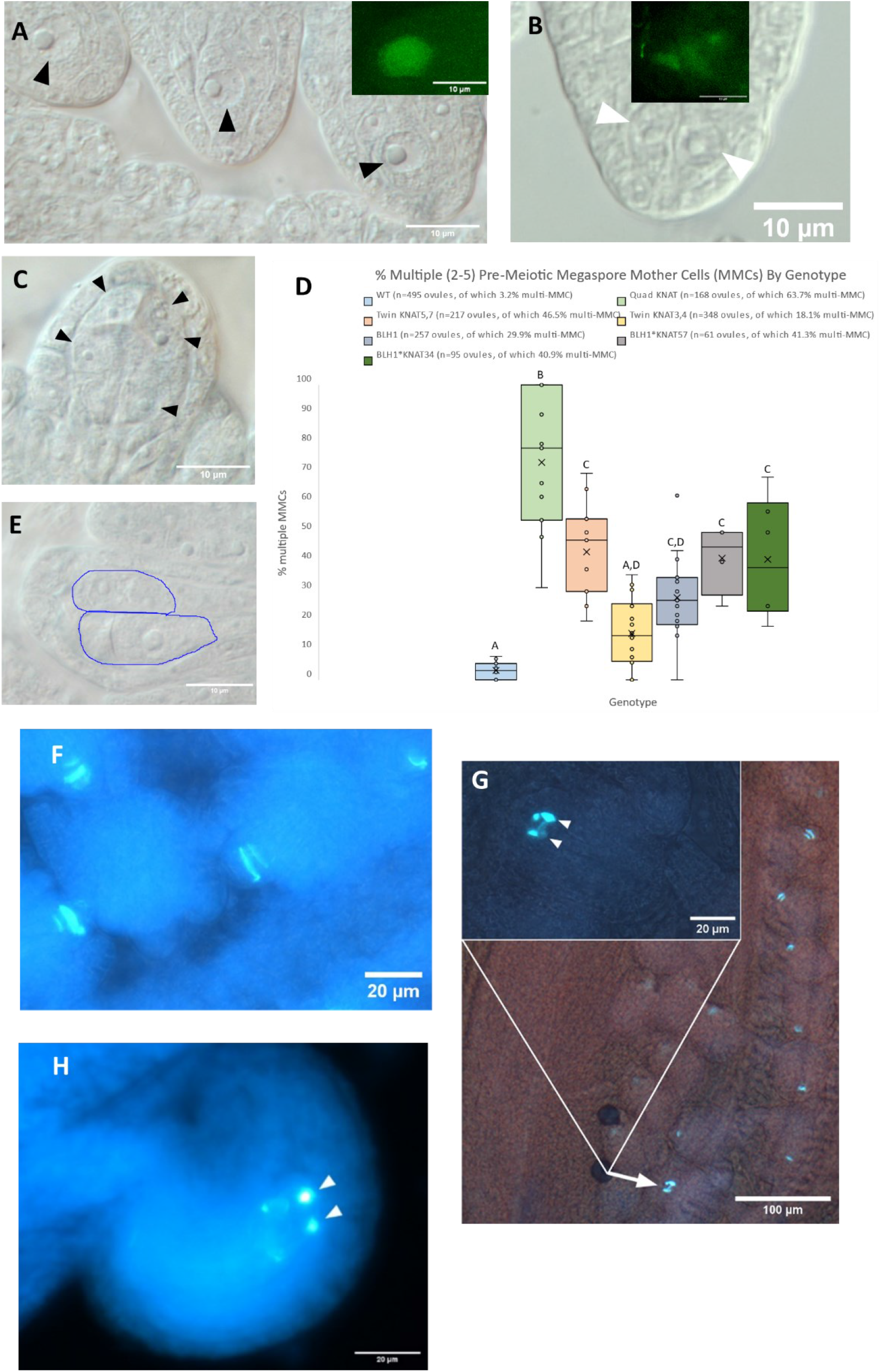
TALE-HD Misexpression results in the specification of multiple sporophytic germlines. **(A)** WT Megaspore other Cells (MMCs) with DIC optics and KNU:YFP confocal false colour image inset. **(B)** duplicate MMCs in KNOX2 misexpression line with KNU:YFP confocal false colour image inset **(C)** Quad KNAT multi-MMC with large number of additional MMCs (indicated by arrows) with bulged nucellus visible. **(D)** quantification of multiple-MMC frequency by genotype. Letters indicate adjusted p-values for p<0.05 from ANOVA and post-hoc Tukey test. **(E)** multiple FG2 stage gametophytes, in *SPL:BLH1* line, indicated with blue outlines **(F) to (H)** aniline blue staining for callose in female meiosis in **(F)** WT and **(G)** *KNAT5,7* expressing ovules, one of which appears to have multiple meiotic cells shown by inset, and clearly has excess callose relative to other ovules. **(I)** *BLH1* expressing ovule showing double post-meiotic gametophytes. Arrows indicate multiple meiotic or post-meiotic cells within a single ovule.

### BLH1 and Class 2 KNOX Expression in the Male Gametophyte has Differing Impacts on Sperm Identity

Unstained pollen nuclei are difficult to visualise, so lines were crossed into the *HTR10:RFP* sperm cell marker. This, combined with DAPI staining and further crossing into vegetative cell marker *VCK:GFP*, allowed the identification of an unusual low-penetrance monospermic pollen phenotype. Sperm identity was confirmed by presence of HTR10, despite there only being a single nucleus present. Although in single Z-sections or projected confocal stacks only a single sperm may be visible (fig 3a, cyan arrows), this can be clarified with inspection of optical sections or 3D-projection. The frequency of monospermic pollen grains (counted using DAPI staining) was 5-10% in homozygous Quad*KNAT*s and twin*KNAT5,7s*, and 0-5% in twin*KNAT3,4s*. It was never observed in wild type. In heterozygous *promSPL:BLH1* plants, this ‘monospermy’ occurred between 4-10%; however these single *promSPL:BLH1* ‘sperm*’* nuclei do not express *HTR10:RFP* expression and thus have failed to acquire full sperm identity. *KNAT34*BLH* lines have a frequency of 6-16%, *KNAT57*BLH* 8-13%, and *QuadKNAT*BLH* 16-20% ‘monospermy’ although it is unclear whether the *BLH*KNAT* pollen grains express HTR10. In the male gametophyte, both *KNATs* and *BLH1* would thus seem to inhibit pollen mitosis 2, but with differing fates conferred on the aberrant sperm/generative cell. *KNATs* could cause an early acquisition of sperm identity or allow sperm differentiation to proceed after DNA replication but before division, while *BLH1* prevents pollen mitosis 2 without allowing sperm differentiation.

**Figure 3.**
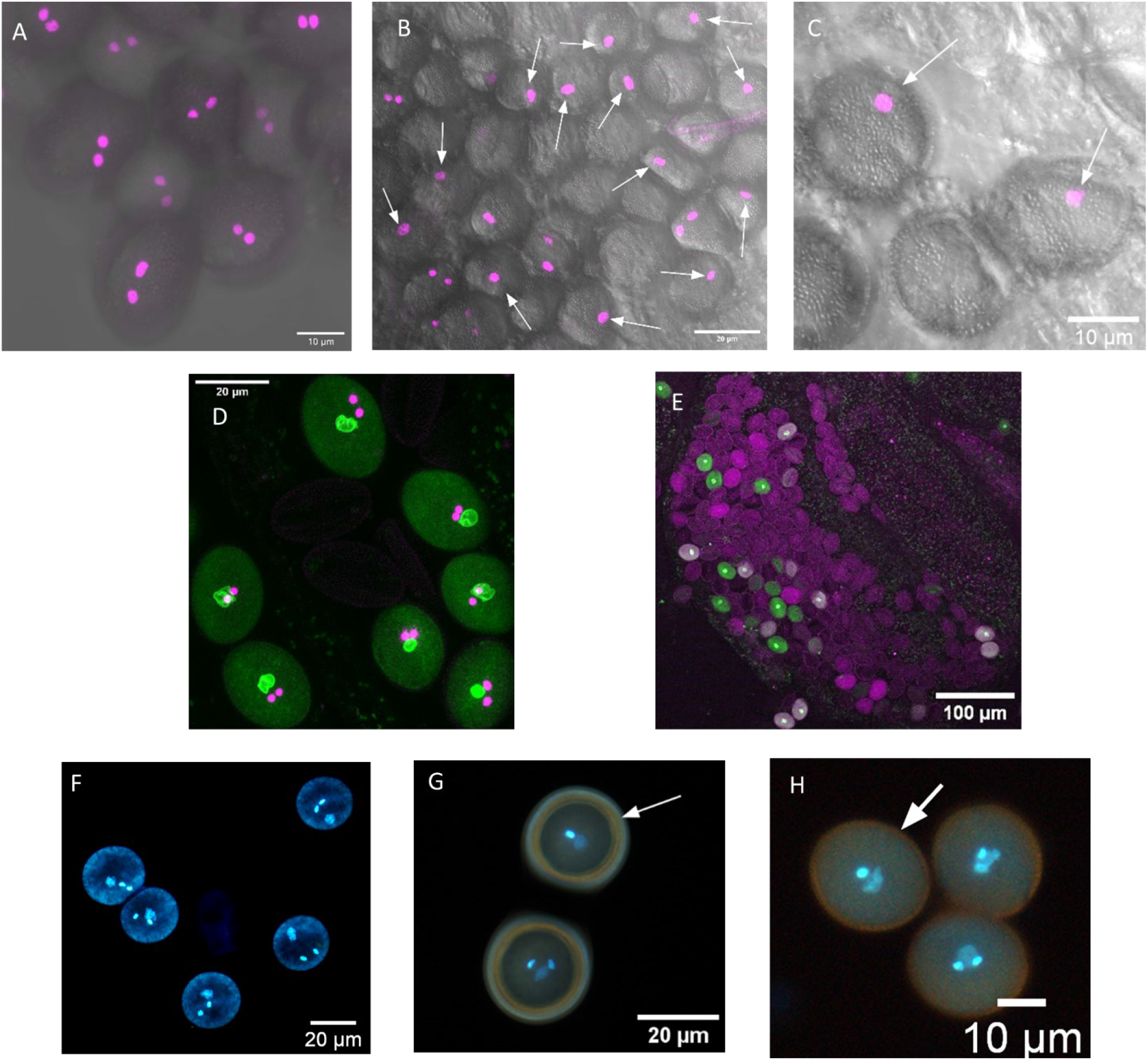
*KNAT* or *BLH1* misexpression inhibits pollen mitosis 2 with differing effects on sperm differentiation. **(A)** R10-RFP Expression in WT pollen grains. **(B) and (C)** HTR10-RFP in pollen grains with KNAT5,7 expression, true monospermic pollen grains shown with white arrows. **(D)** VCK1-GFP and HTR10-RFP expression in WT pollen grains showing vegetative and sperm cell nuclei. **(E)** mostly aborted pollen grains following BLH1 expression, live pollen grains show VCK1-GFP signal in green. **(F)** to **(H)** DAPI stained pollen in **(F)** WT, **(G)** KNAT57, and **(H)** BLH1, with monospermic pollen grains indicated by arrows in (G) and (H).

DAPI staining and the *VCK1:NLS-GFP* marker revealed no abnormalities in the vegetative cell nucleus in *KNAT* expression lines. In *SPL:BLH1*, both *VCK1:NLS-GFP* and *HTR10:RFP* showed reduced expression due to pollen lethality and frequent pollen abortion prior to pollen mitosis 2 (fig3.F).

### Misexpression of *BLH1* or/and *KNAT* in the Female Gametophyte Recapitulates the *Eostre-1*

#### Phenotype

Pagnussat *et al*. (2007) report defects in nuclear migration and cell identity due to gametophytic misexpression of *BLH1* in the *eostre-1* mutant. We observed similar defects as previously reported in the *eostre* mutant (Pagnussat *et al*., 2007) with comparable phenotypes visible in *promSPL:BLH1*, suggesting expression of this construct largely recapitulates *eostre-1* whilst *KNAT* expression does not appear to have much of an effect. FG4 proved to be a key stage of embryo sac development for identification of an ‘*eostre*-like’ fate based on organisation of nuclei pairs (fig.4A-E). Careful inspection allowed the *eostre* phenotype to be identified in mature embryo sacs (fig4.G-H).

**Figure 4.**
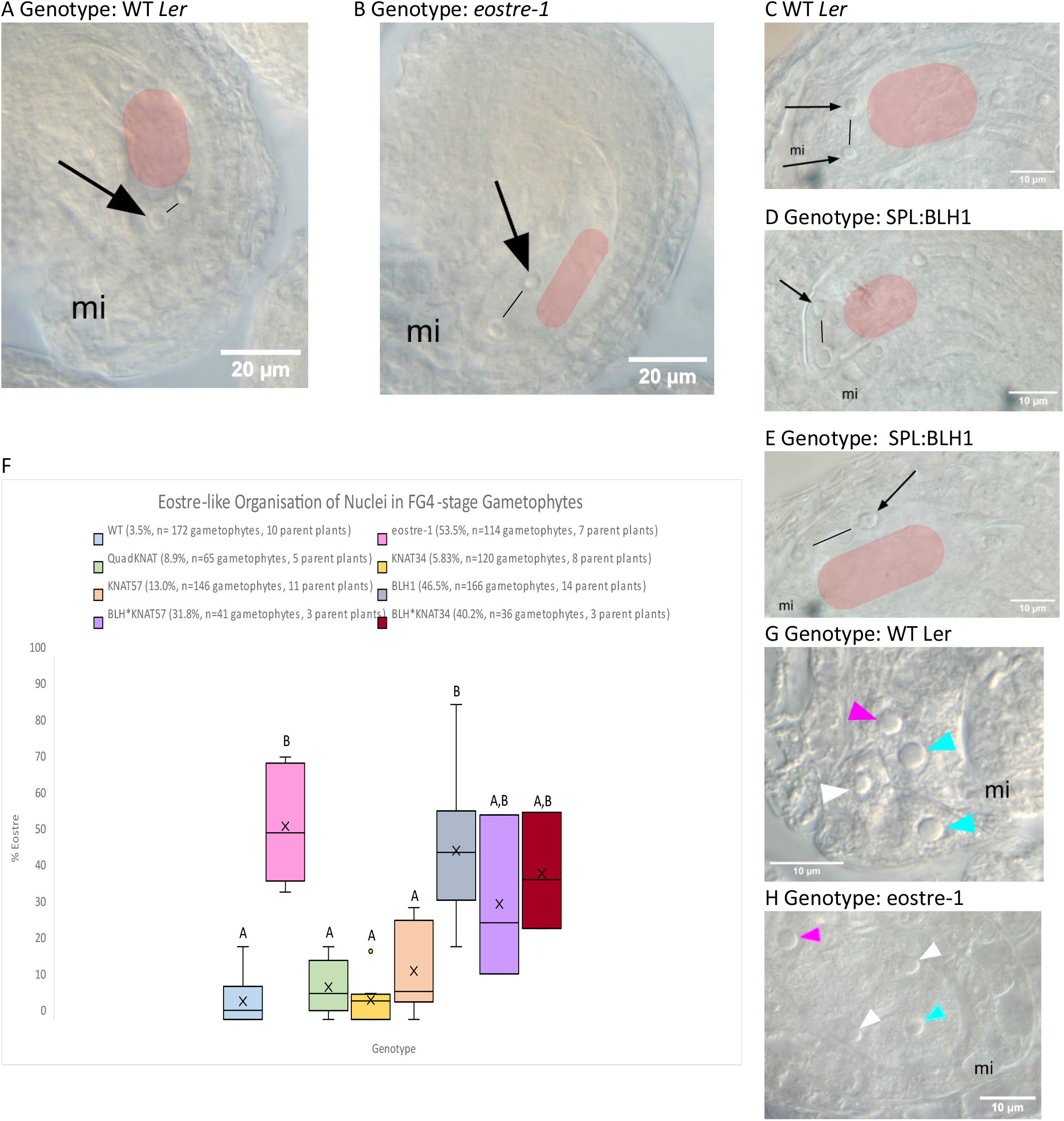
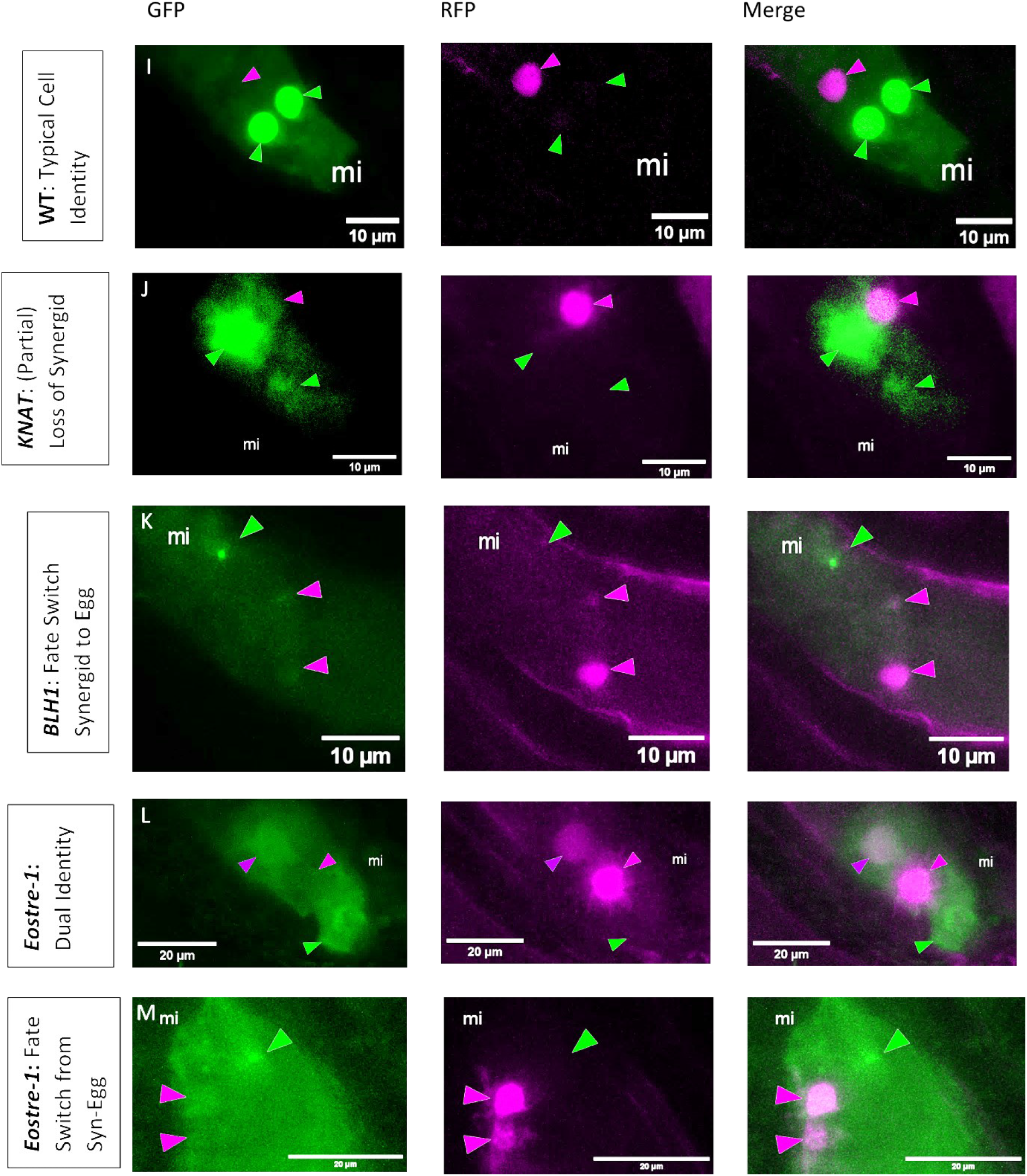
Misexpression of either BLH or KNOX2 recapitulates nuclear migration defects described in the *eostre m*utant. **(A)** to **(E)** FG4 stage female gametophytes in different genotypes with the *eostre* perturbation shown in **(B)** and **(E)** whilst **(A)** and **(C)** show the typical WT FG4 orientation. *pSPL:BLH1* ovules display a mixture of WT **(D)** and *eostre*-like **(E)** phenotypes. Location of the central vacuole in the embryo sac is shown with transparent red block, and black line indicates the angle between the two nuclei at the micropylar end of the embryo sac. The *eostre* perturbation is characterised by this angle being parallel to the vacuole rather than perpendicular as is typical of WT. **(F)** quantification the penetrance of this *eostre* phenotype. **(G)** and **(H)** mature female gametophytes in **(G)** WT orientation and **(H)** with the *eostre* fate switch from synergid to egg. Cyan arrows indicate synergids, white arrows indicate egg cells, and magenta arrows indicate central cells. **(I)** to **(M)** false colour confocal images of the FGR7 embryo sac nuclear identity marker in different genetic backgrounds showing induction of the synergid to egg identity switch by TALE-HD misexpression. Arrows point to synergid (green) and egg cells (magenta) across channels, coloured to reflect the cell type indicated.

**Figure 5.**
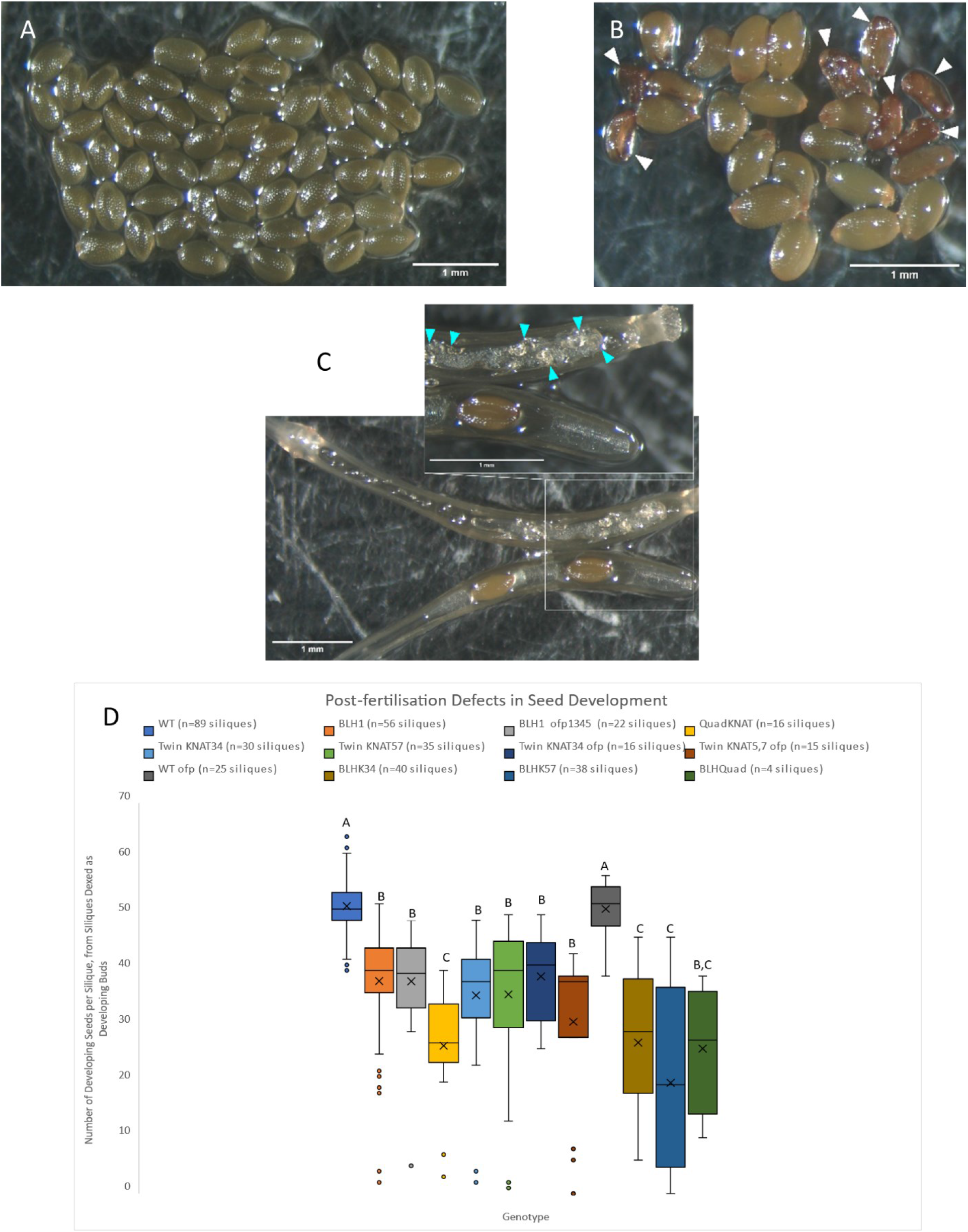
SPL:TALE-HD Expression results in post-fertilisation defects in seeds with a model for the impacts of extopic TALE-HD expression in male and female germline development. **(A)** the complete contents from a near-mature dexed WT silique, **(B)** the complete contents from a near-mature dexed *BLH1* silique, with defective seeds indicated by arrowheads **(C)** a particularly severely impacted *BLH1* silique with aborted ovules indicated by cyan arrowheads. **(D)** quantification of seeds per silique (including defective seeds in mutants) in dexed near-mature siliques.

The FGR7 marker (Volz *et al*., 2013) is a cell identity marker for the mature embryo sac, with Central Cell marked by *NLS-YFP*, synergid *NLS-3xGFP*, and egg *NLS-RFP*. This line therefore provides a molecular indication of identity, complementing the cytology. Our data show that an ‘*eostre*-like’ cell-fate-switch also occurs with *KNAT* misexpression, matching *promSPL:BLH1* and *eostre-1* (fig4.I-N). The FGR7 marker also suggests degrees of fate switch rather than the binary switch visible morphologically. Interestingly, it has been shown (also using this FGR7 marker) that egg cell identity can be conferred by position within the gametophyte (Sun et al., 2021). It remains unclear whether TALE-HD expression impacts cell identity upstream or downstream of nuclear migration.

#### Pre-fertilisation TALE-HD Misexpression Impacts Seed Development

Although *prom:SPL* is not expressed post-fertilisation, lasting seed development defects occur in siliques where *SPL:TALE-HD* was induced pre-fertilisation. This includes malformed seeds and absence of embryos in some mature or nearly-mature seeds that appeared flattened and translucent. There is a significant decrease in seed set in dexamethasone-induced siliques comparing WT and *SPL:BLH1*, or any combination of WT and *SPL:KNAT*. Surprisingly, combining these constructs in the *ofp1345* quadruple-knockout background does not cause a further decrease in seed set compared to WT, suggesting perhaps that more subtle perturbation of the system is needed to detect enhancement of the phenotype, or a minimal role of these *OFP* genes.

## Discussion

### Misexpression of TALE-HDs in Arabidopsis shows a Range of Reproductive Phenotypes

The data and observations from the lines reported here show a surprisingly diverse range of reproductive phenotypes. This study does not attempt to explore any endogenous roles TALE-HDs may play in reproductive development (if any), instead they show that their misexpression is able to disrupt key reproductive processes. Importantly, it can change cell fate in gametophytes and, most strikingly, cell identity in meiocytes and gametes. Whether some, or all, of these changes in cell fate and developmental trajectory can be mapped directly onto the transition from sporophyte to gametophyte remains to be determined, but our data constitute unambiguous evidence that TALE-HD misexpression exerts severe effects on development around the transition between ontogenies, and at fate specification of the sporophytic germline (meiosis) and that of the gametophyte germline (gametogenesis). These results are summarised diagrammatically in Figure 6.

**Figure 6.**
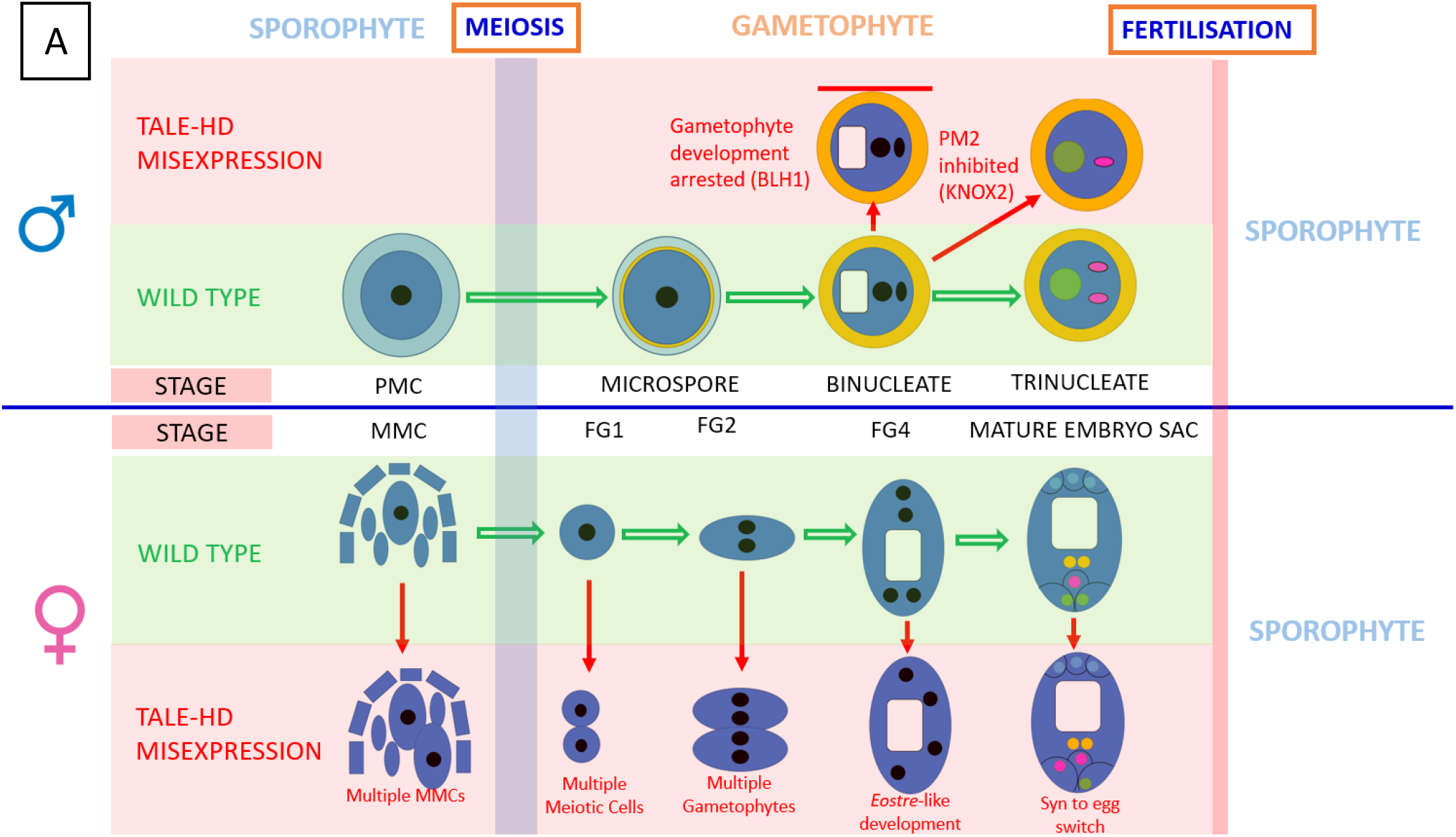
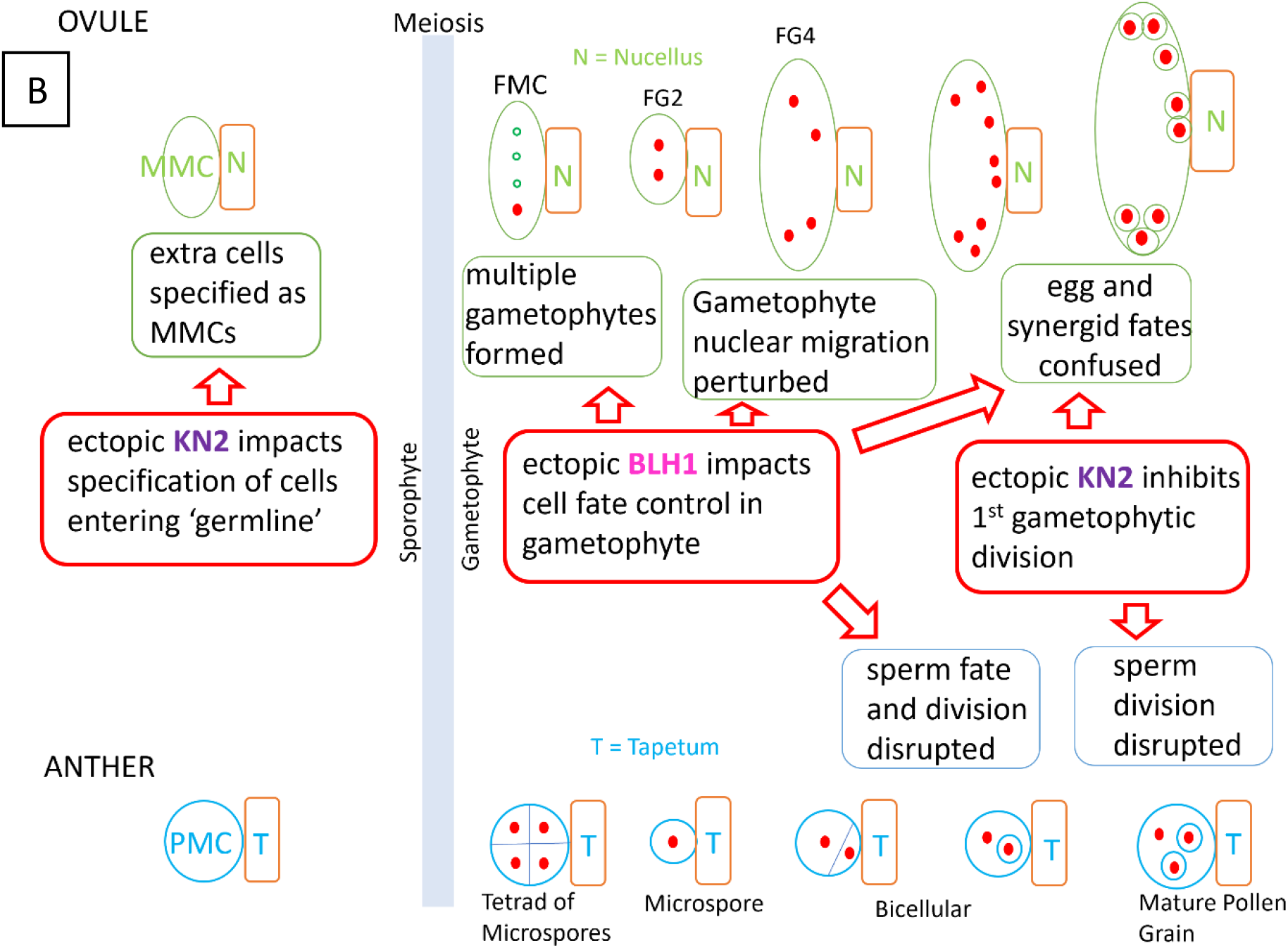
Diagrammatic representations of the effects of ectopic expression of TALE-HD genes in *Arabidopsis* reproductive cell lineages. **(A)** Male development from meiotic cell designation to fertilisation is shown above, female below. WT development is shown in green, misexpression phenotypes described here are shown in red. Nuclei are indicated with black or coloured circles, colours in mature embryo sac and trinucleate pollen reflect the marker lines used. Pale boxes indicate vacuoles, while dark blue indicates cytoplasm. This emphasises the combined functions of BELL and KNOX. **(B)** Presents a similar representation of development, instead emphasising the differing functions of BLH1 and Class 2 KNOX (KN2) in the male and female reproductive lineages.

Unexpectedly, most of our experiments do not reveal a clear difference between phenotypes resulting from ectopic expression of *promSPL:KNAT34* or *promSPL:KNAT57*. Certainly, the lower expression of *promSPL:KNAT34* (fig1) seems to be reflected in lower penetrance of mutant phenotypes, but the similarity of the phenotypes observed suggests the identity of individual KNATs is not crucial in this context. Data from Furumizu et al. (2016) suggest that the specificity of the KNOX-BELL interaction varies between contexts and whether playing endogenous or ectopic roles. Experiments addressing this question, including expressing each KNAT independently, are clearly required. Interestingly, the phenotypes resulting from *prom:SPL:BLH1* expression and those from *KNAT* expression seem similar before and around meiosis (fig2) whilst *BLH1* expression has a much clearer effect on gametophyte development itself (fig4) matching previous reports (Pagnussat *et al*., 2007). Gamete identity defects resulting from *BLH* or *KNAT* expression appear similar for female gametes (fig4) whilst they differ for male gametes (fig3).

### Ectopic TALE-HD Expression Impacts Sporophyte Germline Specification

The MMC can be regarded as the sporophyte female germline, and its specification is under hormonal control, with roles for brassinosteroids (Cai et al., 2022), auxin (Su et al., 2017), and gibberellin (Ferreira et al., 2018). Recent work by Cai et al. (2022) suggests that MMC fate is the default in the pre-meiotic nucellus, which becomes repressed in all but a single, specified cell. The multi-MMC phenotype described here from ectopic KNAT expression may, therefore, involve downstream dysregulation of hormone or epigenetic pathways (Rodríguez-Leal et al., 2015).

It is notable that the multi-MMC frequencies observed here are among the highest reported in the literature (Cai et al., 2022) and there are often more than two MMCs, each of which appears to be able to undergo meiosis and start development as a gametophyte. Interestingly, in many apomictic angiosperm species, including *Hieracium* and *Hypericum*, where ectopic BLH1 expression is known to be important for apomixis (Okada et al., 2013; Galla et al., 2019), atypical development begins from the aposporous MMC with subsequent atypical callose deposition (Galla et al., 2011). Qualitatively, many of the abnormalities observed here following KNAT misexpression appear similar to cytological observations of apospory in these species (Galla et al., 2011) suggesting that *promSPL:KNAT* expression is able to promote some aspects of apomictic development. It has also been suggested that BELL+KNOX homologues in *Chlamydomonas* may activate meiosis (Lee et al., 2008; Harrison et al., 2010) suggesting that this ability for BELL+KNOX to promote a form of aposporous apomixis may be deeply conserved and partly recapitulated in the phenotypes described here.

### TALE-HD TF Expression Most Severely Impacts Gamete Specification

The promotion of egg cell fate by both *promSPL:BLH1* and *promSPL:KNOX2s* shown here matches and extends previous observations (Pagnussat *et al*., 2007), and reveals that *KNAT* expression can also cause this switch. This not only supports the hypothesis that KNOX2 genes are the relevant heterodimersiation partners in this case, but also shows that other KNOX2 genes (in addition to the *KNAT3* shown previously [Pagnussat *et al*., 2007]) can cause this phenotype. Furthermore, the data from our marker line studies (fig 4) provide new evidence that the molecular identity of these nuclei matches the cytological observations. Typically, the absence of the partner TALE-HD for heterodimerisation restricts the severity of ectopic expression phenotypes (Cheong, 2019). Similar to other studies of ectopic KNOX expression in different contexts (Furumizu et al, 2016; Cheong, 2019), combined *BLH+KNOX2* ectopic expression in here seems to enhance each individual phenotype as indicated by the penetrance reported. This effect is probably underestimated in these data, as ovules expressing both frequently aborted around meiosis, suggesting that coexpression can be gametophytic lethal.

We show that *promSPL:TALE-HD* expression, of either *BLH1* or *KNAT*, promotes female gamete (egg) identity, whilst *KNAT* promotes, or permits, male gamete (sperm) identity which *BLH1* seems to inhibit. This particularly severe effect of misexpression on gametes is comparable to findings from *Marchantia* and *Chlamydomonas* and supports the suggestion by Hisanaga *et. al* (2021) that aspects of the sporophyte control function in *Physcomitrium* are derived with a conserved role around gametogenesis and meiosis. The results described here support this hypothesis, perhaps over the Horst and Reski (2016) model. Distinguishing between impacts on gametophyte development and on gametogenesis is important when interpreting these findings, due to ambiguity in the literature and to ensure the homologous developmental stage is being studied in each taxon when examining conservation of the mechanisms for controlling the switch in ontogeny.

The *promSPL:KNOX2* monospermic pollen phenotype is particularly striking, and despite apparent failure of pollen mitosis 2 HTR10 expression nonetheless occurs, indicating either premature acquisition of sperm cell identity or that sperm cell differentiation is somehow able to proceed despite the generative cell not having divided. *HTR10* is a direct target of *DUO1* (Borg et al., 2011), the key master regulator of sperm cell identity (Rotman et al., 2005) which is conserved across plants (Higo et al., 2018). The *duo1* mutant (Rotman et al., 2005) shows some phenotypic similarity to the observed monospermy, however in *duo1 HTR10* expression does not occur and sperms never differentiate, making this a failure in sperm identity (Borg et al., 2011) rather than early acquisition. Instead, *Duo1* is more similar to *promSPL:BLH1* expression, where acquisition of sperm cell identity is inhibited with a failure of pollen mitosis 2. Some kind of sex-specific phenotype is not necessarily unexpected, as it does also occur in *Marchantia* (Dierschke *et al*., 2022; Hisanaga *et al*., 2022)

## Conclusions

Inducible Golden Gate constructs designed to ectopically express *KNAT*s and *BLH1* in the male and female germlines of *Arabidopsis* have been successfully generated and validated, thus generating a resource that allows, for the first time, the testing for a conserved ability for BELL/KNOX to trigger apomictic transitions in the alternation of generations, as has been shown for bryophytes (Horst *et al*., 2016; Horst and Reski, 2016). The approach of using BpiI digestion of cDNA to distinguish between expression of a ‘domesticated’ olden Gate transgene and the endogenous copy (fig1) proved a useful addition to the standard Golden Gate pipeline. Inclusion of a blind mutation in a suitable restriction site in all such constructs to permit this strategy would be useful addition to standard methodology.

This work extends the previously neglected findings of Pagnussat *et al*. (2007), making use of modern molecular marker tools to provide further clarity on gametophytic expression of *BLH1* including in male development. The results also further support a *KNAT*+*BLH1* role in this phenotype. Class 2 KNOX genes are relatively little-studied (Furumizu et al., 2015); here we show that – as is the case for *BLH1* - repression of KNOX2 is crucial for gametophyte development and germline specification in flowering plants. This work also represents the first explicit test of whether the Horst and Reski (2016) model of a *Physcomitrium* sporophyte control module is conserved in land plants for controlling alternation of generations; results are summarised in fig6. In angiosperms, meiosis and gametogenesis are separated by only two or three mitotic divisions, making it difficult to separate the distinct processes of germline specification, control of meiosis, gametophyte development, and gametogenesis. The results presented here suggest that *KNAT*+*BLH1* misexpression exerts a surprisingly substantial effect on meiosis and in gametogenesis. Further work to identify what exactly is being dysregulated by *promSPL:TALE-HD* expression is now needed to improve our understanding of cell-fate control in plant reproduction and investigate whether this system could be employed to manipulate plant reproduction in agriculture.

## Methods

### Plant Care and Handling

All *Arabidopsis* plants were grown in controlled environment chambers (Gallenkamp) at the Department of Plant Sciences, University of Oxford, at 18.5°C in 16h light/8h dark. Individual lines were combined by crossing with presence of inserts verified by PCR.

Induction of transgenes uses the dexamethasone inducible LhGR-pOp6 system (Craft *et al*., 2005). Induction ‘dexing’ was performed following Schubert et al (2022) with the same treatment applied to experimental and WT lines in all cases. GUS staining (Jefferson et al., 1987) was performed on harvested inflorescences in 24 well plates.

### Generation of Plant Lines

All constructs were generated using Golden Gate (GG) cloning (Engler et al., 2013). *In silico* design was carried out using Benchling and Snapgene software. Golden Gate plasmids were transformed into *Arabidopsis* through floral dip (Clough and Bent, 1998) using the FAST red selection marker (Shimada et al, 2010) for identifying transformants and homozygotes.

### RT-qPCR

RNA extraction was performed using the QIA R asy Plant Mini Kit (manufacturer’s protocol. R A from plants with 4+ inflorescence apices were used; pre-fertilisation buds from two inflorescences were snap frozen in liquid nitrogen and further inflorescences from the same plant were labelled, ‘dexed’, and snap frozen 18 hours after induction. cDNA was synthesised using qPCRBIO cDNA Synthesis Kit (PCR Biosystems) and qPCR was performed in 10 μl reactions using qPCRBIO SyGreen Mix (PCR Biosystems) with a StepOnePlus Real-time PCR system (Applied Biosystems). For verifying transgene induction, total expression of each *KNAT* or *BLH1* were compared in pre- and post-induction samples from the same plant. This does not distinguish the endogenous gene and averages expression across the tissue. To distinguish the transgene, cDNA was digested using BpiI (Thermofisher Scientific) since, for Golden Gate cloning, BpiI sites were removed. Thus, digestion cleaves the endogenous cDNA but not the domesticated gene. Subsequent qPCR using primers around the cleavage site is then specific to construct-driven expression. All qPCR data are from 4-6 biological replicates and two (averaged) technical replicates.

### Microscopy

For brightfield microscopy, inflorescences were fixed in FAA fixative (3.7% formaldehyde, 15% acetic acid, 50% ethanol) in 24-well plates for 12-16h, transferred to 30% ethanol for ∼15mins, washed with 30% ethanol, and transferred to lactic acid-glycerol clearing solution (20% lactic acid, 20% glycerol, in 1xPBS) for 4-48 hours. Post-fertilisation tissues were cleared using chloral hydrate clearing solution (8:2:1 chloral hydrate:water:glycerol).

For DAPI staining of mature pollen, 3-4 open flowers were placed in 300μl DAPI staining solution (100mM NaH_2_PO_4_, 1mM EDTA, 0.1% Triton X-100, 0.4 μg/ml DAPI) and vortexed. Flowers were removed leaving pollen in solution, which was pelleted by two-minute centrifugation and mounted on a slide (protocol provided by Michael Borg, *pers comm* 2022). For earlier stages of pollen development, buds of appropriate stage were crushed on a slide with a brass rod in DAPI solution and visualised after 20min incubation. Aniline Blue staining for meiotic callose was performed as described in Cai *et al*., (2022) and samples mounted in 30% glycerol 1% DABCO antifade in 1xPBS. Non-fluorescence light-microscopy was performed using a Zeiss Axiophot with Chromyx HD camera, using brightfield or Nomarski DIC illumination. UV microscopy for DAPI and Aniline Blue was performed on an Olympus BX-50 with mercury vapour lamp and captured using a QImaging QICamFast1394 in QCapture Pro. Size was calibrated via stage micrometer. Confocal microscopy was performed on a Leica SP5 or Zeiss 880 upright confocal laser scanning microscope. DAPI and Aniline Blue were excited at 405nm, YFP and GFP at 488nm, and RFP at 561nm.

### Data Analysis

Images were processed in the FIJI distribution (Schindelin et al., 2012) of ImageJ (Schneider et al., 2012) for processing of confocal stacks, cropping, addition of annotations and scale bars, and optimising brightness/contrast. Micrographs were not otherwise modified and adjustment to brightness/contrast were uniform across the image. Multi-MMC data, *eostre* phenotype penetrance, and seed count data were analysed by ANOVA and Tukey-HSD post-hoc test using R (R Core Team, 2017). RT-qPCR data were analysed via ΔΔCt using Thermo isher DataConnect Design&Analysis software.

## Acknowledgements

We thank Saher Mehdi (formerly of Department of Plant Sciences, University of Oxford) for early help with molecular biology, Daniela Vlad (Department of Biology, University of Oxford) for advice on Golden Gate Cloning, and Vinay Shukla (Department of Biology, University of Oxford) for help with qPCR. Pollen marker lines were kindly provided by Michael Borg (Max Planck Institute for Biology, Tubingen) and the FGR7 embryo sac marker by Rita Groß-Hardt (University of Bremen). The KNU:YFP line was generated by Matthew Tucker (University of Adelaide) and the *ofp* knockout lines by Yu Sun (Hebei Normal University).

## Notes

### Competing Interest Statement

The authors have declared no competing interest.

